# A collagen orientation switch reshapes fin architecture

**DOI:** 10.64898/2026.01.17.700086

**Authors:** Rintaro Tanimoto, Kazuhide Miyamoto, Koji Tamura, Shigeru Kondo, Junpei Kuroda

## Abstract

The orientation and distribution of fibrillar collagen are critical determinants of the shape and mechanical properties of bones and organs.^1–3^ However, how they are spatially organized within tissues is still poorly understood,^4–7^ as visualizing these collagen architectures remains challenging. Actinotrichia (AT), the spear-shaped fibrillar collagen structures located at the distal tips of fish fins, are easily observable due to their large size and distinctive morphology^8–14^ and have recently emerged as a model system for studying collagen fiber organization.^15–19^ In this study, we generated knockout lines for the fish-specific extracellular matrix (ECM) genes *actinodin1* and *actinodin2* (*and1/2*), which are lost in tetrapods.^12^ Loss of these genes dramatically altered the orientation of collagen fibers, thereby inducing changes in fin morphology. In the wild-type fins, AT are orderly arranged beneath the epidermis, forming layers parallel to the fin surface, and their individual fibers radiate distally toward the fin tip. In contrast, double knockout (dKO) of *and1/2* results in overall fin reduction accompanied by increased thickness. Examination of the collagen structure distribution revealed the presence of aberrant collagen fibers oriented perpendicular to the fin epidermis. Moreover, the vertically oriented fibers contributed to thickening of the mesenchymal region in which they were distributed. The number of abnormal fibers increased with the severity of *and1/2* deficiency, suggesting that collagen fibers in fins inherently tend to align perpendicular to the epidermis when these genes are absent. Furthermore, in tetrapods lacking the *and* gene family—specifically amphibians, the tetrapod group most closely related to fish^20^—examination of the developing limb, the organ homologous to paired fins in fish,^21^ revealed collagen fibers oriented perpendicular to the epidermis. The distribution pattern also resembled that observed in the fin buds of *and1/2* dKO fish. Together, these findings highlight collagen patterning alterations as a previously unrecognized factor contributing to the evolutionary divergence between thinned fins and thickened limbs. Moreover, the identification of mutants that dramatically alter collagen fiber orientation is unprecedented, suggesting that analysis of Actinodin (And) function unveil the mechanisms underlying collagen matrix formation.^22–28^

## RESULTS

### Loss of *and1/2* reorients collagen fibers and alters fin morphology in zebrafish larvae

A previous report analyzed the function of And1/2 by simultaneously knocking *and1/2* down with morpholinos.^12^ However, this approach might only partially suppress the function of these proteins. Therefore, in this study, we generated a dKO zebrafish line of *and1/2* using CRISPR-Cas9^29^ (Figures S1A and S1B). First, we examined the phenotype of the caudal fin fold because AT is distributed throughout this tissue and can be readily observed.^9,11^ Specifically, we performed high-resolution 3D imaging of collagen structure within the fin fold using our previously reported diaminofluorescein-FM diacetate (DAF) staining method^19,30^ and analyzed the relationship between collagen fiber orientation and tissue morphological changes. Figures 1A and 1B show fluorescence images of the caudal fin fold captured from a lateral view. Compared with the wild-type, the dKO exhibited a reduced fin size, with lengths shortened along both the proximodistal and dorsoventral axes (Figure 1C). Each magnified fluorescence images of the area surrounding the anterior end of the notochord is shown in the panels below (Figures 1A’ and 1B’). In the wild-type, the AT radiated distally from the base of the caudal fin fold. Their thickness were uniform (2–3 µm) and equivalent to that of typical collagen fibers (1–20 µm) found in vertebrate tissues.^22^ In contrast, in the dKO, the overall fluorescence of the fin tissue was reduced, and the radially oriented fibers were thinner than wild-type AT (Figure 1B’, white asterisk). Furthermore, particularly in the region near the base of the caudal fin fold, strong punctate fluorescence was observed (Figure 1B’, white arrowhead). Therefore, to clarify the structural features of this fluorescence signal, we enlarged the relevant region and performed a 3D reconstruction (Figures 1D and 1E). In the wild-type, fibrous structures were aligned immediately beneath the epidermis on both sides, with no collagen-like structures visible in the spaces between them. In contrast, in the dKO, fibers running parallel to the epidermis were present, but these fibers were thinner than wild-type AT (Figure 1E, white asterisks). Moreover, aberrant fiber bundles appeared in the interstitial spaces (Figure 1E, white arrowhead), which were not present in the wild-type. Measurement of the angles of these abnormal fibers relative to the epidermis revealed that their distribution was not random but clearly biased toward a perpendicular orientation (Figure 1F). Abnormal bundle thickness and spacing were largely constant, with maxima of ∼3 µm. Consequently, imaging from a lateral view showed that the signals exhibited a spotted pattern (Figure 1E, white arrowhead). In addition, optical section images revealed that the thickness of the fin interstitial space in the dKO increased by 20% compared with the wild-type (Figures 1D’, 1E’, and 1G).

**Figure 1.**
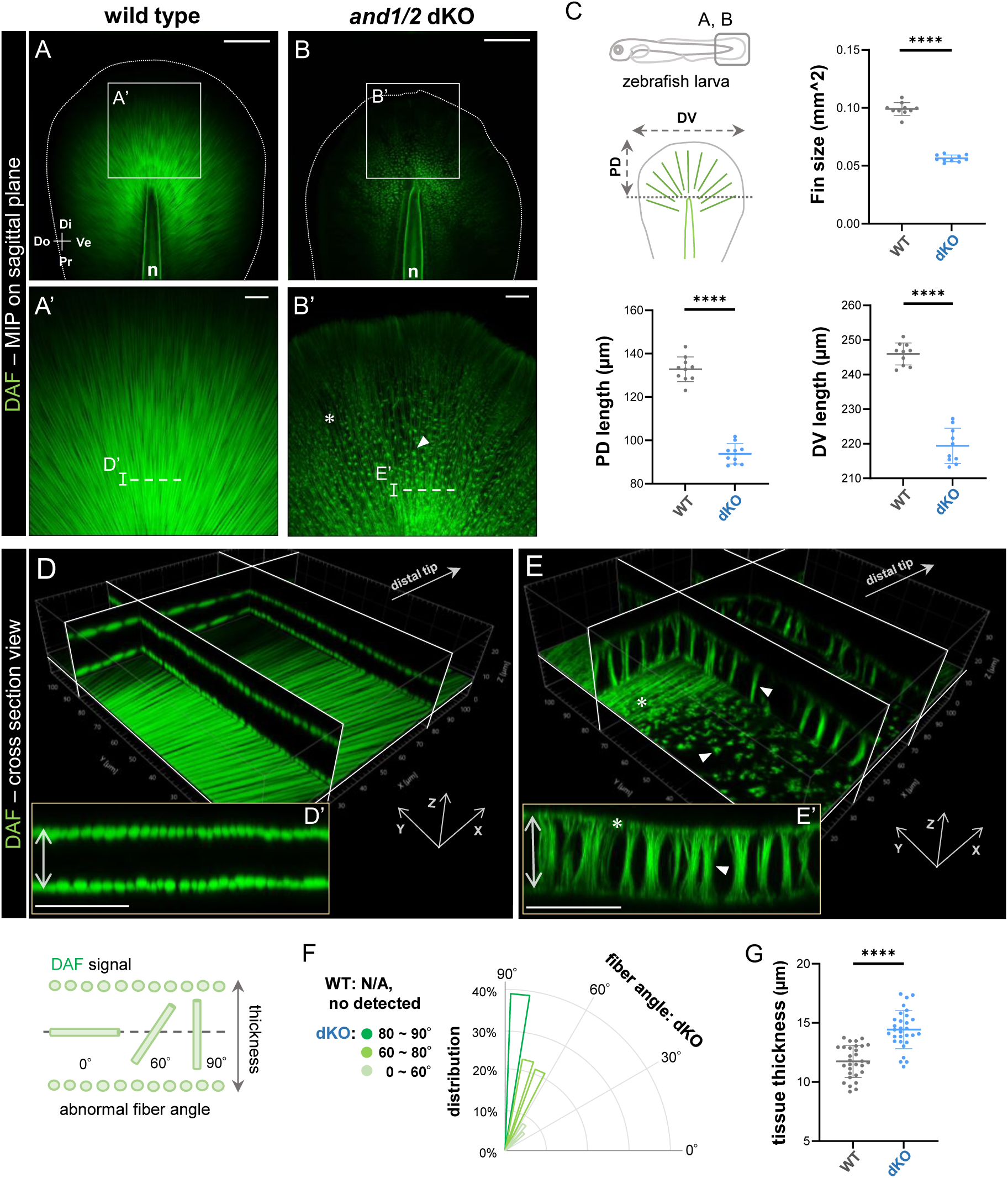
Double knockout of *and1/2* alters the organized alignment of collagen fibers in the caudal fin fold of zebrafish. (A-B’) Confocal images of the caudal fin fold in wild-type and *and1/2* dKO zebrafish at 5 days post-fertilization (dpf). Collagen fibers were fluorescently labeled (green) by whole-mount DAF staining. Magnified views of the white-boxed regions are shown in the lower panels. White dotted lines: fin outlines; white asterisk: abnormally thin fibrous structures; white arrowhead: abnormal particle-like structures. Pr: proximal; Di: distal; Do: dorsal; Ve: ventral; n: notochord; MIP: maximum intensity projection. Scale bars: 100 µm (A, B), 20 µm (A’, B’). (C) Quantification of fin morphology. Welch’s t-test: **** p<0.0001. WT vs. dKO; N = 10 per group. (D, E) Orthogonal slice view from (A’, B’) (MIP, 5-µm stack (D’, E’)). White asterisks: abnormally thin fibers; white arrowheads: abnormally oriented fibers; white double-headed arrows: thickness of the interstitial space. Scale bars: 20 µm. (F) Distribution of abnormal fiber angle measured from optical section. WT: N/A; dKO: N = 172 (10 larvae per group). (G) Quantification of fin mesenchyme thickness (the region 50 µm from the notochord end). Welch’s t-test: **** p<0.0001. WT vs. dKO; N = 30 per group (10 larvae per group.)

**Figure S1 related to Figure 1.**
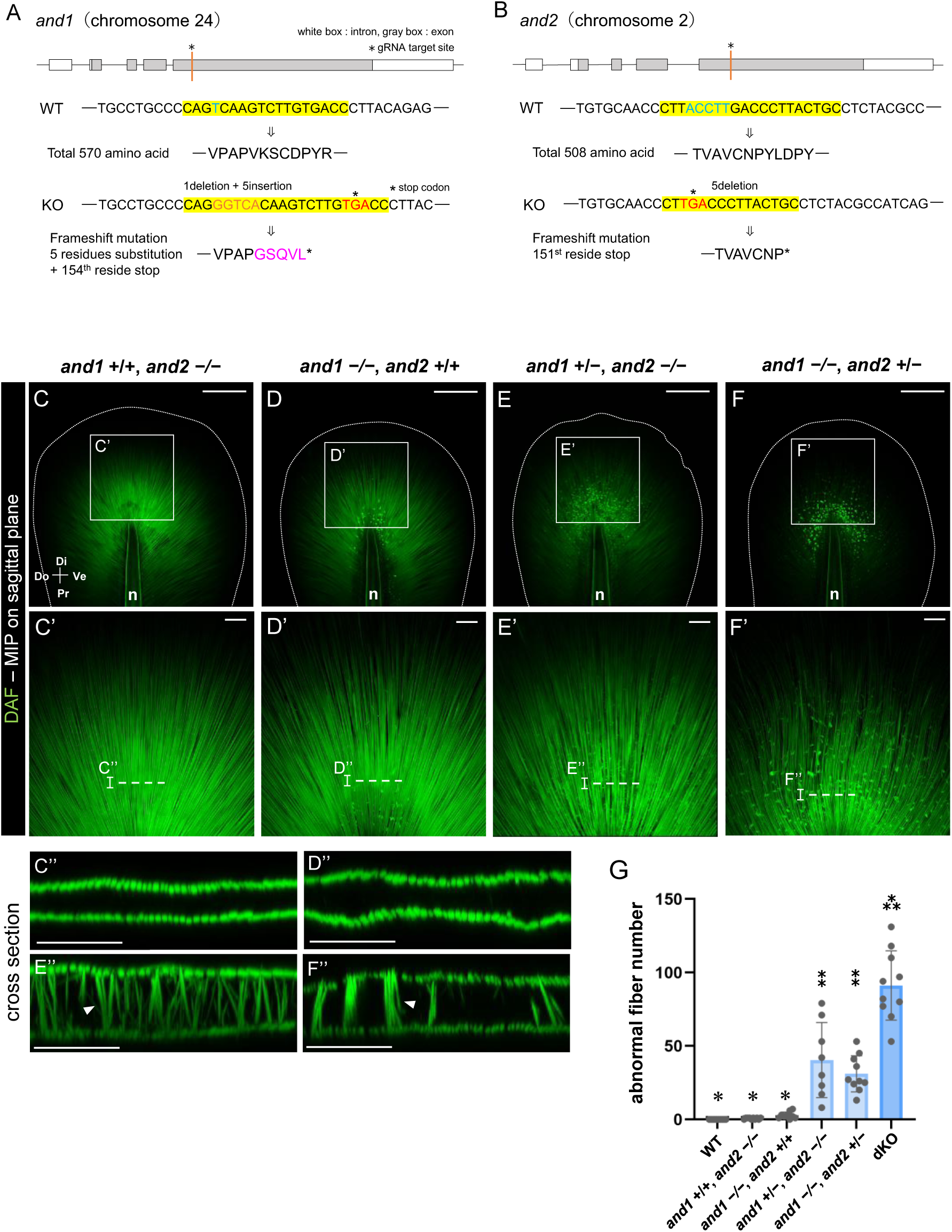
Collagen fiber orientation in the caudal fin folds of zebrafish from different *and1/and2* mutant genotypes. (A, B) Locus information for the wild-type (WT) vs. knockout (KO) alleles of *and1* / *and2*. Frame-shift mutations indicate that each *and1/2* KO sequence lacks more than 70% of the amino acids in the respective WT-encoded proteins. These annotations were referenced from the UCSC Genome Browser (Zebrafish May 2017 release (GRCz11/danRer11), *and1*_mRNA (chr24:26279503-26283359, NM_001197254) / *and2*_mRNA (chr2:31763202-31767652, NM_001111190). Yellow box: gRNA target region; Blue text: Deletion site; Orange text: Insertion site; Magenta text: Substituted amino acid. (C-F’’) Confocal images of caudal fin folds from each *and1* and *and2* knockout genotype at 5 days post-fertilization (dpf). Collagen fibers were fluorescently labeled (green) by whole-mount DAF staining. Magnified views of the white-boxed regions are shown in the lower panels. Cross-sectional views (MIP, 5-µm stack) corresponding to panels C’-F’ are also shown below. White dotted lines: fin outlines; white arrowheads: abnormal vertical fibers. Pr: proximal; Di: distal; Do: dorsal; Ve: ventral; n: notochord; MIP: maximum intensity projection. Scale bars: 100 µm (C-F), 20 µm (C’-F’’). (G) Quantification of the number of abnormal vertical fibers (the region 50 µm from the notochord end). Tukey-Kramer method; different symbols indicate p<0.05. N (left to right: WT to dKO) = 10, 10, 9, 8, 10, 10.

**Figure S2 related to Figure 1.**
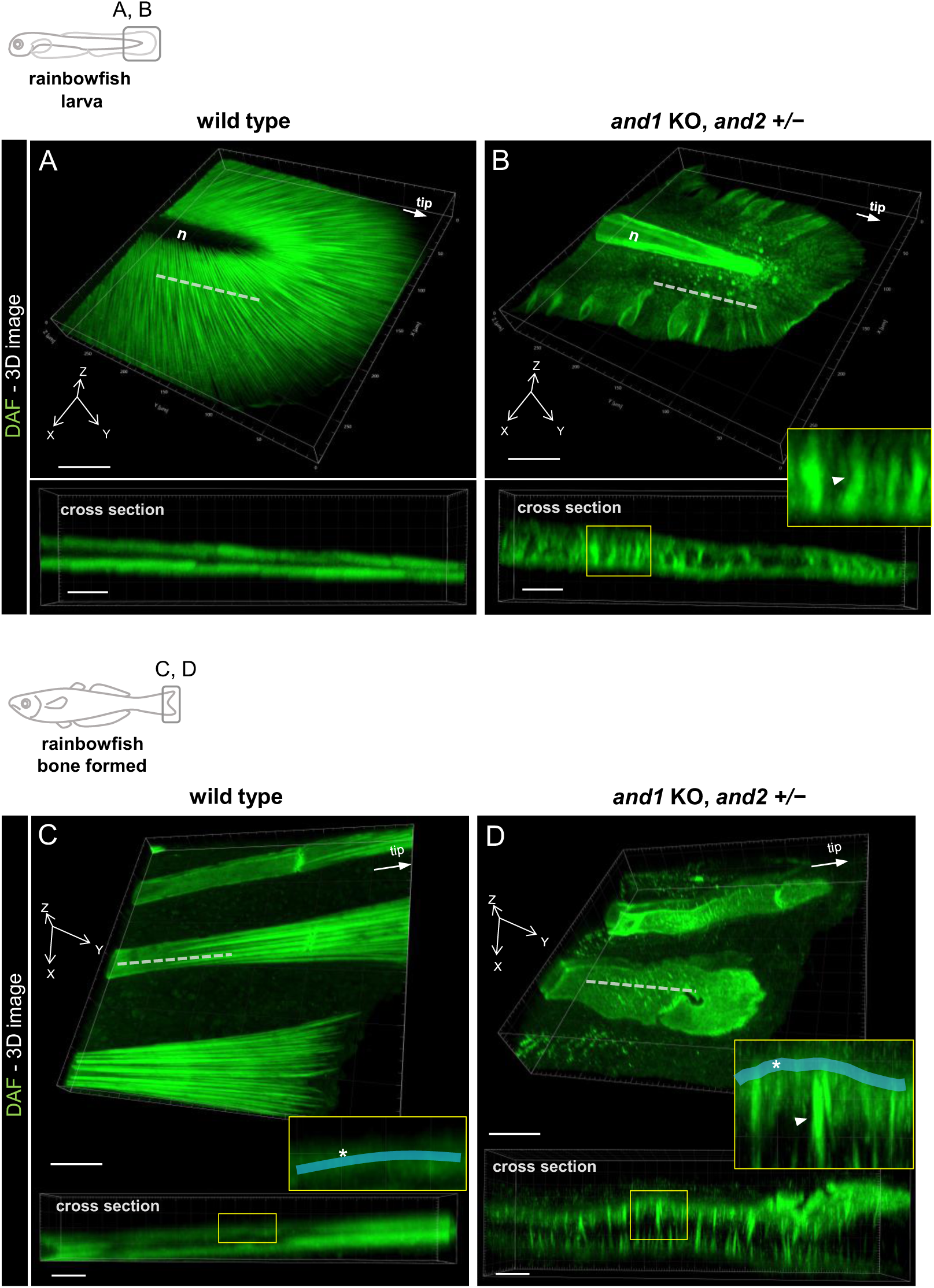
Abnormal vertical fibers are distributed in the caudal fin of *and1/2* double mutant rainbowfish, resembling that in the *and1/2* dKO zebrafish line. (A, B) Confocal images of the caudal fin fold in wild-type and *and1/2* double mutant rainbowfish at 7 dpf. Collagen fibers were fluorescently labeled (green) by whole-mount DAF staining. Cross-sectional views at white dotted lines are displayed on lower panels. (C, D) Confocal images of the caudal fin tip in wild-type and *and1/2* double mutant rainbowfish at DF-st4. Collagen fibers were fluorescently labeled (green) by whole-mount DAF staining. Cross-sectional views at white dotted lines are displayed on lower panels. White arrowheads: abnormally oriented collagen structures; white asterisk with translucent blue lines: collagen matrices within fin bones. Scale bars: 50 µm (3D views, A-D), 10 µm (cross-sectional views, A-D).

Next, to determine the contribution of each gene to this phenomenon—the formation of abnormally oriented fibers—we similarly observed the collagen structures in larvae of each genotype: *and1 +/+*; *and2 -/-, and1 -/-*; *and2 +/+, and1 -/+*; *and2 -/-, and1 -/-*; *and2 -/+*. In the first two genotypes, no significant changes were observed in the distribution of collagen matrix (Figures S1C and S1D). In the latter two genotypes, abnormal fibrous structures were observed similar to the dKO (Figures S1E” and S1F”, white arrowhead), but their number was significantly lower compared with the dKO (Figures S1E and S1F). The relationship between the number of abnormal fibers and each genotype is shown in Figure S1G. Thus, *and1* and *and2* are presumed to share the same function in collagen orientation, namely directing collagen fibers to align parallel to the fin epidermis. Conversely, in the absence of *and* genes, these collagen fibers shift to a perpendicular orientation relative to the fin epidermis. Furthermore, because the phenotype observed in the zebrafish *and1/2* dKO lines was unprecedented, we performed the same observations using DAF staining on the recently reported *and* mutant of rainbowfish^31^ to determine whether this phenomenon is specific to zebrafish or shared among fish. As shown in Figures S2A and S2B, we confirmed the presence of vertical collagen structures around the base of the caudal fin fold of the mutant, similar to those observed in zebrafish, whereas such structures were absent in the wild-type.

### A shift in collagen orientation reorganizes mesenchymal cell layers

Next, since the orientation of collagen fibers is likely determined by surrounding cells,^9,17,18,32,33^ we examined the state of these cells by fluorescently labeling their nuclei (Figures 2A–2C). Figures 2A and 2B show optical section images of a coronal cross-section of the caudal fin fold (same field of view as Figure 1D’). No obvious differences were observed in the nuclei of epidermal cell (ECs) forming the tissue outer layer between the wild-type and the dKO. ECs in both genotypes exhibited a flattened morphology along with the fin plane. In contrast, significant changes were observed in the distribution and morphology of the nuclei of mesenchymal cells (MCs) within mesenchyme. In the wild-type, the MCs were located immediately inside of the AT (Figure 2A’). However, in the dKO, the MCs were positioned between the vertical fibers (Figure 2B’, white arrowhead), rather than immediately beneath the fibers distributed on the basal side of the epidermis (Figure 2B’, white asterisk). The angular distribution of the major axis of MCs (approximated as ellipses) relative to the epidermis is shown in Figure 2C. This analysis confirms that in the wild-type, almost all MCs were flattened along the AT, whereas in the dKO, they tended to be oriented perpendicular to the epidermis. Additionally, we performed comparative observations of caudal fin fold sections using transmission electron microscopy (TEM) to examine in detail abnormalities in collagen structures and alterations in cell distribution (Figures 2D and 2E). In the wild-type, the epidermal cells (ECs), the AT, and the mesenchymal cells (MCs) were arranged in distinct layers from the outer to the inner side of the tissue, forming a stacked structure (Figure 2D). High-magnification imaging of the boundary region between these three layers revealed that the AT, approximately 2–3 µm in diameter, were orderly aligned immediately beneath the ECs. In addition, characteristic crystalline structures previously reported for AT were clearly observed within them (Figure 2D′).^9,18,34^ The MCs formed a layer on the inner side of the AT, adhering to their surface, while the mesenchyme further inward was extremely thin. On the other hand, in the dKO, although the overall layered organization of the tissue was comparable to that of the wild-type, the mesenchyme displayed a clearly different appearance (Figure 2E). In the region where the AT is present in the wild-type, collagen structures with diameters of several hundred nanometers were scattered and aligned parallel to the epidermal layer (Figures 2E and 2E’, green asterisks). The average width of their distribution was approximately 3 µm, consistent with the width of the AT layer in the wild-type. Moreover, the mesenchyme further inward from this region was clearly thicker than that in the wild-type. The MCs were scattered throughout this area, and surrounding them were vertically oriented collagen bundles, each several hundred nm thick (Figures 2E and 2E′, green arrowheads). A characteristic striped pattern was visible within these structures, consistent with the typical 67-nm pitch of collagen fibers.^1,22^ These observations supported the findings from confocal imaging using DAF staining. Together, they indicate that loss of And function causes the orientation of collagen fibers in the mesenchyme to shift from a parallel to a perpendicular relative to the fin epidermis. This might not only orient the mesenchymal cells along the ectopic axis but also drive tissue expansion in the same direction in the proximal region. Consequently, while distal elongation of the larval fin is suppressed, tissue thickness is inferred to increase.

**Figure 2.**
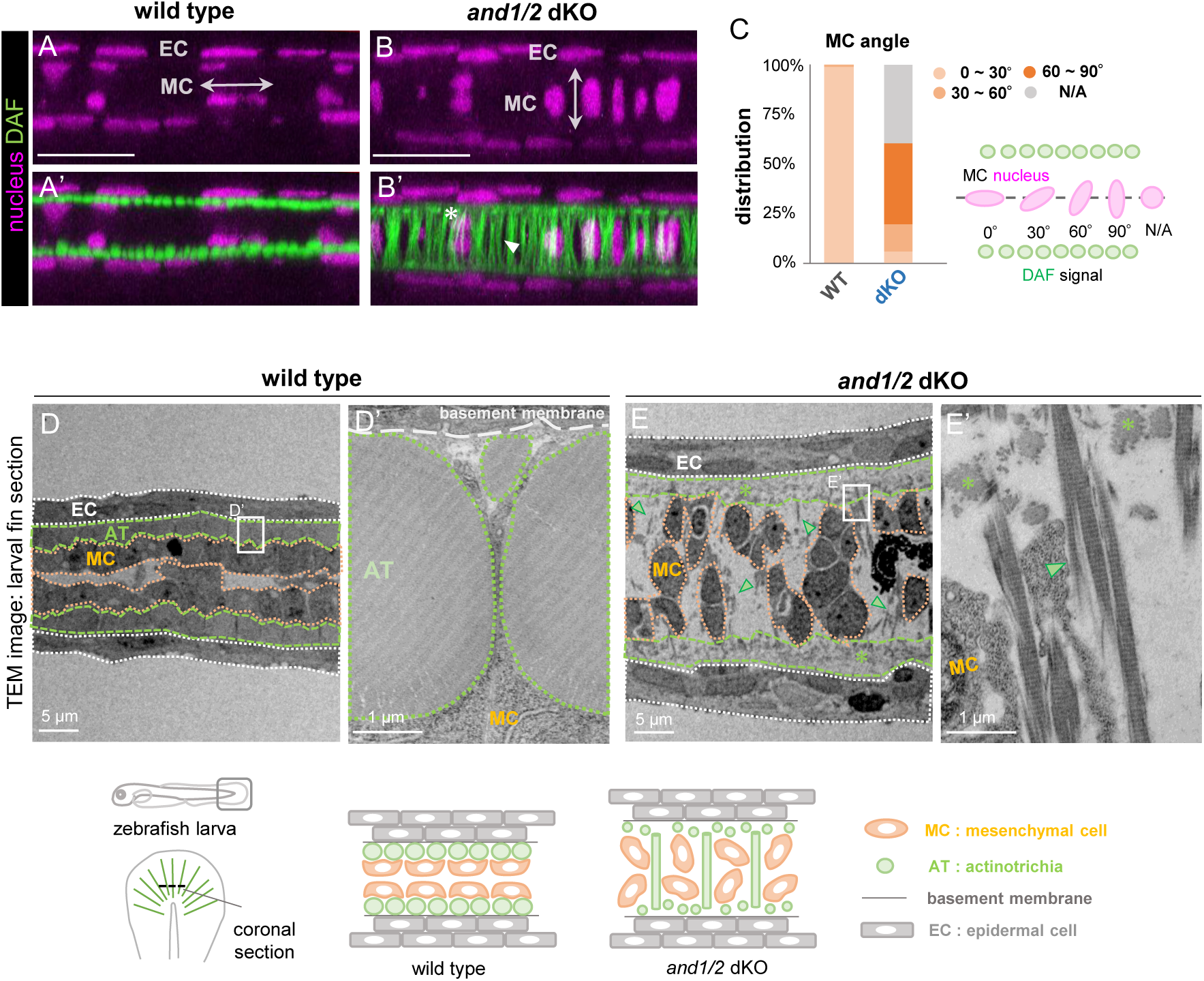
Abnormal mesenchymal cell elongation and collagen fiber orientation in the caudal fin fold of *and1/2* dKO zebrafish. (A-B’) Representative optical sections from wild-type and *and1/2* dKO zebrafish at 5 dpf (MIP, 5-µm stack). Collagen fibers were fluorescently labeled (green) by whole-mount DAF staining and cell nuclei were stained with Hoechst (magenta). White asterisk: parallel-oriented fibers; white arrowhead: perpendicular-oriented fibers. MC: mesenchymal cell nucleus; EC: epidermal cell nucleus. Scale bars: 20 µm. (C) Distribution of mesenchymal cell nuclear orientation relative to the sagittal line. WT: N = 92; dKO: N = 106 (9 larvae per group). (D-E’) TEM images for the coronal section of caudal fin fold in wild-type and *and1/2* dKO zebrafish at 8 dpf. Green asterisks: parallel collagen fibers; green arrowheads: perpendicular collagen fibers. Schematic diagram of larval fin tissue anatomy is shown in the lower panel.

### Alterations in collagen orientation and fin morphology in the *and1/2* dKO persist even after bone formation

During the larval stage described above, AT are present throughout the fin. By contrast, in adults, fin rigidity is provided by fin bones, and AT are restricted to the growing tips of these bones.^8,10,13,14^ To determine whether collagen orientation changes occur under this condition, we next performed a detailed analysis of the caudal fin phenotype in the *and1/2* dKO line. Accordingly, to test whether the collagen abnormalities seen in larva also occur at the fin tips in adult, we performed DAF staining and examined their distribution (Figures 3A and 3B). Figures 3A and 3B show fluorescence images of the fin tip tissue imaged from a lateral view. In the wild-type, the fin bones connected by segments extended linearly along the distal axis, forming regularly spaced rectangular units (Figure 3A, white asterisk). Furthermore, strong fibrous fluorescence signals aligned along this axis were detected in the terminal ∼ 200 µm region of each fin bone. As shown in the magnified image, the AT were densely distributed at the bone tip (Figure 3A’). In cross-sectional views of the bone tip (Figure 3A”), the AT were arranged parallel to the inner surface of the bone. However, in the dKO, the rectangular fin-bone units were distorted, and no fibrous structures were observed at the bone tip (Figure 3B’). The distance from the fin-bone tip to the distal edge of the fin tissue along the proximodistal axis was significantly reduced (Figures 3A–3C). Remarkably, cross-sectional observation of the distal tip region of the fin bone clarified abundant collagen structures running perpendicular to the bone (Figure 3B”). In addition, the mesenchymal tissue filling the space between the left and right fin bones were markedly thickened (Figures 3A”, 3B”, and 3D). Hence, the fin tip tissue of the adult also exhibited changes in collagen orientation and tissue morphology, similar to those observed in the larval stage. These alterations were further supported by TEM observations, as shown in Figures 3E and 3F. Specifically, in the wild-type, the AT were observed immediately inside the fin bone (Figures 3E and 3E’, blue region), but in the dKO, no AT-like structures were detected in this region. Instead, collagen fibers arranged perpendicular to the fin bones were observed, forming a bundled pattern similar to that seen in the larval fin (Figure 3F’, green arrowhead). The mesenchymal tissue between the left and right fin bone layers was also significantly thicker than that in the wild-type (Figures 3E and 3F). Consistently, this phenomenon—collagen fibers oriented vertical relative to the fin bones due to *and* deficiency—was also observed in post-ossification rainbowfish (Figures S2C and S2D).

**Figure 3.**
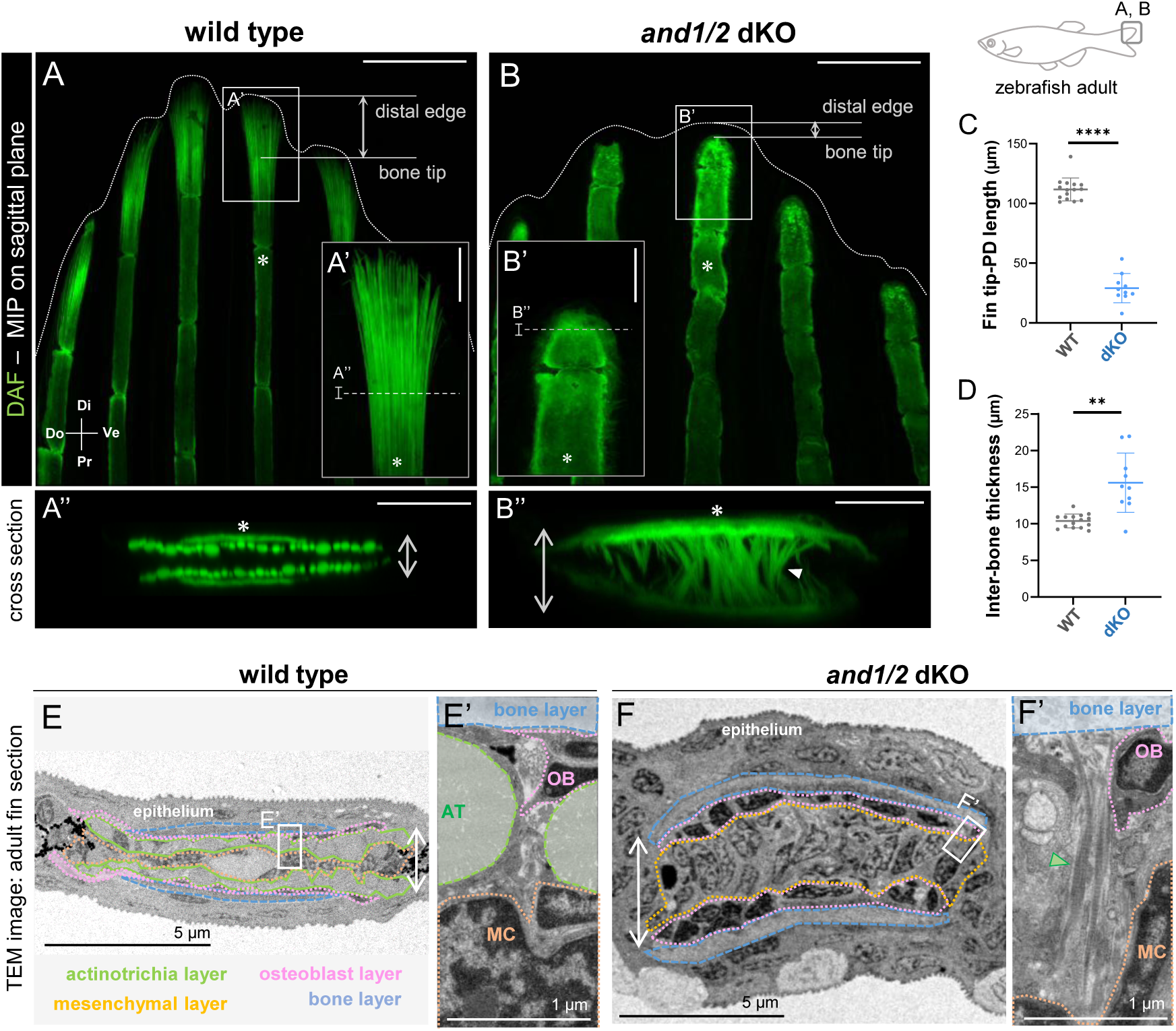
Abnormal collagen fiber distribution in the caudal fin of *and1/2* dKO zebrafish. (A-B’’) Confocal images of the caudal fin tip in wild-type and *and1/2* dKO zebrafish with standard length (SL) = 2.1 ± 0.1 cm. Collagen fibers were fluorescently labeled (green) by whole-mount DAF staining. The magnified images in the white box are shown as insets. Cross-sectional views at white dotted lines are shown on the lower panels (MIP, 10-µm stack). White dotted lines: fin outlines; white asterisks: collagen matrices within fin bones; white arrowhead: abnormally oriented collagen structures; white double-headed arrows: thickness between left and right fin bones. Scale bars: 200 µm (A, B), 50 µm (A’, B’), 20 µm (A’’, B’’). (C-D) Quantification of fin bone morphology. Welch’s t-test: ** p < 0.01, **** p < 0.0001. WT: N = 14; dKO: N = 10 (10 fishes per group). (E-F’) TEM images for the coronal section of caudal fin tip in wild-type and *and1/2* dKO zebrafish with SL = 2.1 ± 0.1 cm. Green arrowhead: abnormally oriented collagen fiber; white double-headed arrows: thickness between left and right fin bones. AT: actinotrichia; MC: mesenchymal cell; OB: osteoblast.

**Figure S3 related Figure 3.**
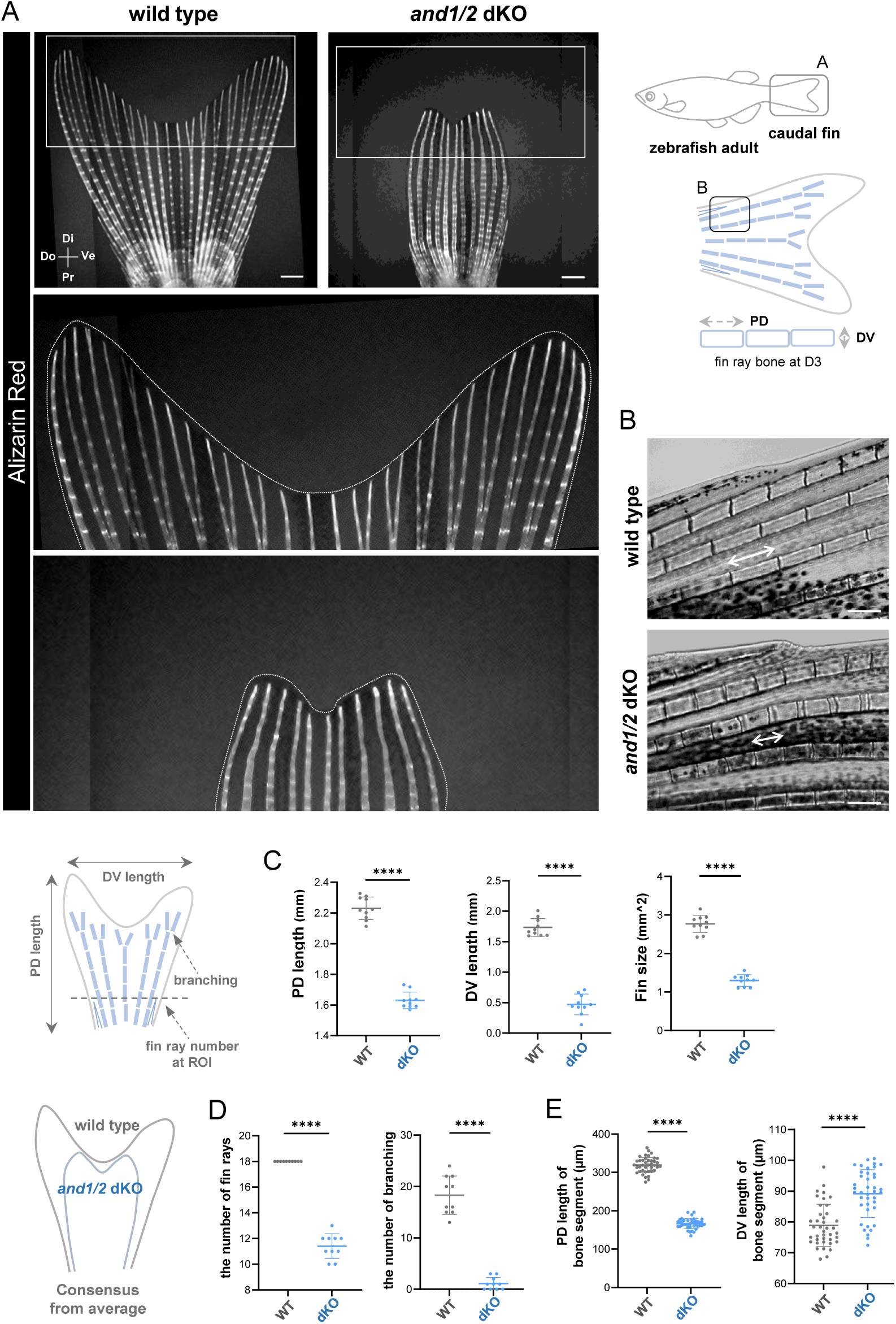
*and1/2* dKO zebrafish exhibit morphological defects in the caudal fin. A) Fluorescence images of the caudal fin stained with Alizarin Red in wild-type and *and1/2* dKO zebrafish with standard length (SL) = 2.1 ± 0.1 cm. White dotted lines: fin outlines. (B) Magnified images of the fin bones in wild-type and *and1/2* dKO. Double-headed arrows indicate the single-segment structure of fin bones. Scale bars: 200 µm (A, B). (C-E) Quantification of fin morphology. Welch’s t-test: **** p < 0.0001. WT vs. dKO. Fin skeleton: N = 10 per group; Bone segments: N = 40 per group (10 fishes per group).

**Figure S4 related to Figure 3.**
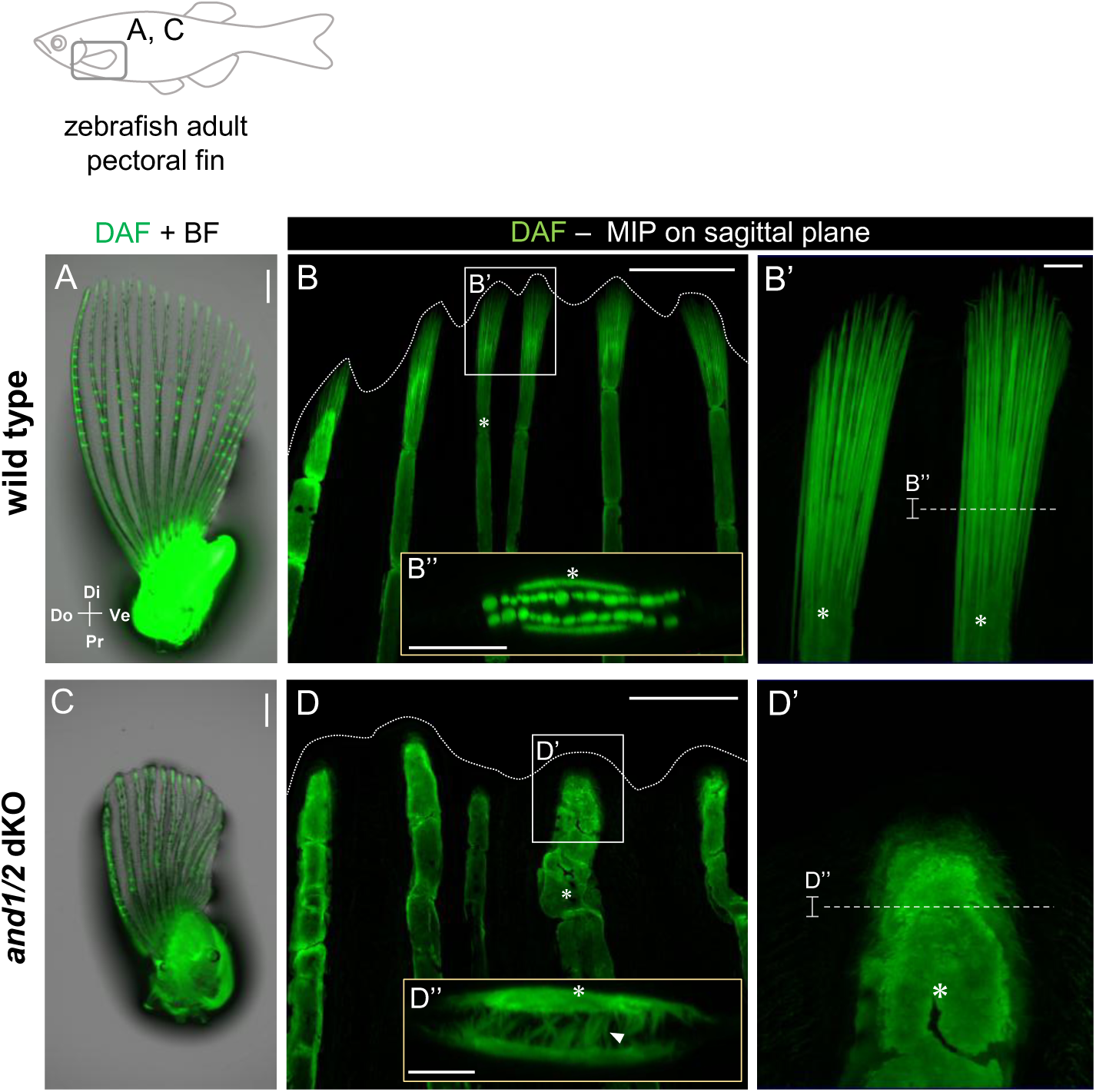
*and1/2* dKO zebrafish display same abnormality in the pectoral fin as in the caudal fin. (A, C) Fluorescence images of the pectoral fin stained with DAF in wild-type and *and1/2* dKO zebrafish with SL = 2.1 ± 0.1 cm. (B-B’’, D-D’’) Confocal images of the pectoral fin tip in wild-type and *and1/2* dKO (MIP, 10-µm stack (B’’, D’’)). White dotted lines: fin outlines; white asterisks: collagen matrices within fin bones; white arrowhead: abnormally oriented collagen structures. Scale bars: 200 µm (A-D), 20 µm (B”, D”).

As shown in Figure 3, the dKO zebrafish also exhibited abnormalities in the fin bones, which form the skeletal framework of fins. Therefore, we also analyzed morphological changes in the caudal fin (Figure S3). This analysis confirmed that the dKO caused fin shortening, abnormal ray patterning, and misshaping of bone segments (Figure S3). The same morphological changes were also observed in the pectoral fins (Figure S4), consistent with a previous study.^35^ Moreover, as in the caudal fin, a similar alteration in collagen distribution was confirmed (Figure S4). These findings indicate that, *and* deficiency induces a common set of phenotypes—altered collagen orientation, local tissue thickening, and distal suppression of fin elongation—that arise regardless of life stage or fin type and are broadly conserved among fish.

### Perpendicular collagen-fiber organization is shared by *and*-deficient fins and limbs

As mentioned above, functional loss of the fish-specific proteins And1/2 caused collagen fibers in the fin to align perpendicular to the epidermis, accompanied by tissue thickening along the direction of collagen orientation. If this finding has evolutionary significance,^12^ collagen fibers should tend to adopt a perpendicular orientation beneath the epidermis of the organ that corresponds to the fin in vertebrates lacking the *and* gene family—that is, in tetrapods. Thus, we examined the distribution of collagen fibers beneath the epidermis in limb buds of amphibians—the tetrapod lineage most closely related to fish^20^—and in the homologous structure,^21^ the paired fin bud, using the same method applied to the caudal fin fold of zebrafish. Figures 4A and 4B show fluorescence images of the collagen fiber organization in the pectoral fin buds of zebrafish. In the wild-type, similar to the caudal fin fold, the AT were neatly aligned only in a direction parallel to the epidermis. In contrast, in the dKO, in addition to collagen fibers aligned in the same direction (Figure 4B’, white asterisk), aberrant collagen fibers were formed and extended across the mesenchyme (Figure 4B’, white arrowhead). As shown in Figure 4C, the angular distribution of these structures revealed a distinct bias toward a perpendicular orientation relative to the epidermis. Moreover, fin bud shortening and increased thickness of the mesenchyme surrounding the endoskeletal disc were also observed (Figures 4A’ and 4B’). Finally, to examine collagen-fiber formation during limb development—the organ homologous to the paired fins—we analyzed fluorescent 3D images obtained from axolotl limb buds using the same method (Figures 4D and 4E). In both forelimb and hindlimb buds, collagen fibers extending into the mesenchyme were observed (Figures 4D and 4E, white arrowhead). Notably, the orientation distribution was indistinguishable from that observed in the *and1/2* dKO fin buds, with both showing approximately 80% of fibers oriented between 70° and 90° (Figures 4C and 4F). Hence, we found that collagen fibers oriented perpendicular to the limb-bud epidermis were present in amphibians, which innately lack the *and* gene family. These findings demonstrate that, in the absence of both And1/2, subepidermal collagen fibers in fin buds and limb buds adopt a perpendicular orientation to the epidermis.

**Figure 4.**
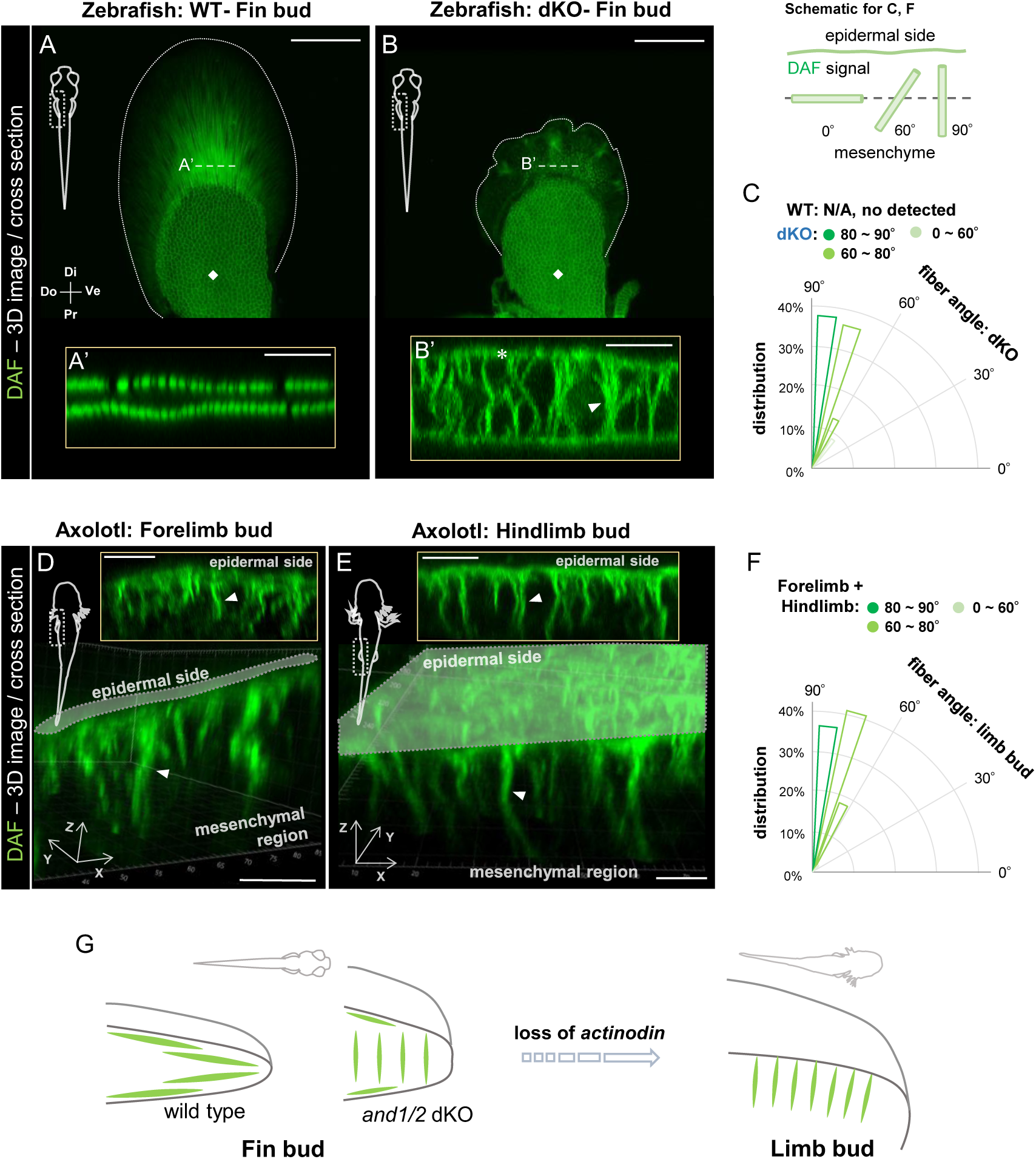
Aberrant collagen fiber orientation arises in the pectoral fin buds of *and1/2* dKO zebrafish, comparable to that observed in axolotl limb buds. (A-B’) Confocal images of the pectoral fin bud in wild-type and *and1/2* dKO zebrafish at 5 dpf (MIP, 5-µm stack (A’, B’)). White dotted lines: fin outlines, white squares: endoskeletal discs, white asterisk: parallel-oriented fiber, white arrowhead: vertically oriented fiber. Scale bars: 100 µm (A, B), 10 µm (A’, B’). (C) Distribution of abnormal fiber angle in pectoral fin bud measured from optical section (the region 10 µm distal to the endoskeletal disc edge.) WT: N/A (8 larvae); dKO: N = 121 (4 larvae). (D-E) Confocal images of limb bud in axolotl. White arrowheads: collagen fibers perpendicular to the epidermal layer. Scale bars: 10 µm. (F) Distribution of fiber angle in limb bud measured from optical section. N = 336. (G) Schematic diagram illustrating the novel finding of this study.

## DISCUSSION

In this report, we revealed that the fish-specific gene *and1/2* is essential for establishing the ordered alignment of collagen fibers in fin tissue and for forming an elongated, thin fin structure. Analysis using high-sensitivity 3D imaging demonstrated that *and1/2* deficiency induces abnormal vertical alignment of collagen fibers, resulting in fin-bud shortening and thickening, and transition to a limb-like morphology. Furthermore, we found that collagen fibers with a similar vertical orientation are also formed in the limb buds of amphibians, a tetrapod group most closely related to fish. Our findings suggest that the acquisition of this collagen pattern represents an important prerequisite for the emergence of limbs.

In fish fins, AT are known to align parallel to the epidermal layer,^8,9^ and previous studies suggested that And1/2 are involved in AT production as AT are composed of these ECM proteins.^12,14,18^ In this study, we generated *and1/2* dKO fish and applied a highly sensitive collagen-fiber imaging method we recently reported.^19,30^ This allowed us to specifically demonstrate a previously overlooked role for And1/2: beyond AT formation, And1/2 function promotes fin-tissue thinning by preventing the vertical alignment of collagen fibers. A future challenge is to elucidate the molecular mechanism by which And1/2 interacts with collagen fibers and controls their orientation pattern.^22,25,36^ This is expected to provide a molecular and cellular understanding of how the unique collagen fiber orientation in fin tissue is generated.^17–19,37^ Furthermore, our findings deepen insight into the evolutionary significance of the loss of the *and* gene family^12,35,38–43^; such a loss may have promoted the vertical alignment of collagen fibers in the ancestor of tetrapods, thereby contributing to the fin-to-limb transition. This represents a novel finding demonstrating that changes in ECM pattern formation can serve as a key driver of morphological evolution, complementing conventional models of cell-cell communication mediated by signaling molecules and transcription factors.^44–48^ In recent years, the ECM has been recognized not merely as a supportive structure but as a dynamic element controlling cell proliferation, differentiation, and migration.^49–51^ Particularly, the formation of fibrillar collagen during the elongation phase of fin and limb buds suggests that changes in the collagen orientation are likely decisive factors in morphogenesis.^52–55^ Collectively, our findings support a potential mechanism whereby And1/2 loss-of-function rearranges collagen-fiber organization and mesenchymal-cell dynamics, thereby converting fin buds into a limb-bud-like configuration.^33,56,57^

Overall, this study proposes that fibrillar-collagen patterning represents a pivotal driver of vertebrate appendage variation.

## ACKNOWLEDGMENTS

We thank Midori Tanaka and Rie Taniguchi, technical staff members at Kondo Laboratory, Graduate School of Frontier Biosciences, Osaka University, for fish maintenance. We also thank Dr. Ritsuko Suyama, Dr. Yasuhiro Hirano, Dr. Ryosuke Kaneko, and Dr. Ritsuko Morita for managing the confocal microscopes at the Graduate School of Frontier Biosciences, Osaka University. We acknowledge the Leica imaging Lab at Osaka University for the support in obtaining fluorescence imaging data. This research was funded by JSPS KAKENHI, Grant Number 20H05943, JSPS KAKENHI, Grant Number 21K06200, JSPS KAKENHI, Grant Number 23H04301, JST FOREST Program, Grant Number JPMJFR224P, and JST SPRING, Grant Number JPMJSP2138.

## AUTHOR CONTRIBUTIONS

Conceptualization, J.K.; methodology, R.T. and J.K.; investigation, R. T., K.M., and J. K.; writing – original draft, R.T. and J.K.; writing – review & editing, R.T., K.M., K.T., S.K., and J.K.; visualization, R.T., J.K.; supervision, J.K.; project administration, J.K.; funding acquisition, S. K., K. T., J.K., and R.T.

## MATERIALS & METHODS

### Animal maintenance

The AB strain was used as the wild-type zebrafish in all experiments. Zebrafish and rainbowfish were maintained under standard conditions at 28 °C with a 14 h light/10 h dark photoperiod. Albino axolotls were maintained at 20 °C under the same 14 h light/10 h dark cycle. Animals were anesthetized with tricaine (MS-222) at concentrations adjusted according to body size. All zebrafish and axolotl experiments were approved by the Animal Care and Use Committee of Osaka University. All rainbowfish experiments were approved by the Animal Care and Use Committee of Tohoku University.

### Establishment of *and1/and2* double knockout (*and1/2* dKO) zebrafish line

To generate founder fish for the *and1/2* dKO line, guide RNAs (gRNAs) targeting *and1* (5′-AGGGTCACAAGACTTGACTGGGG-3′) and *and2* (5′-AGGCAGTAAGGGTCAAGGTAAGG-3′) were designed. The gRNAs and Cas9 protein (NEB, M0646) were co-injected into one-cell–stage fertilized eggs of wild-type AB zebrafish, producing the F0 mosaic generation. Individuals in the F0 population exhibiting abnormal fin morphology were selected and intercrossed to obtain the F1 generation, which also displayed fin abnormalities. F1 individuals were crossed with wild-type fish, and their offspring were genotyped (see Methods section below). Fish carrying knockout alleles for both *and1* and *and2* (Figure S1A, B) were selected as the F2 generation (*and1* −/+, *and2* −/+). Intercrossing the F2 population yielded the actinodin1/actinodin2 double-knockout (*and1/2* dKO) zebrafish line.

### Genotyping of zebrafish

For DNA extraction, whole embryos or half bodies of larvae were collected, and caudal or anal fin samples were collected from adult fish anesthetized with 0.02% tricaine (MS-222; Sigma, A5040) in tank water. Samples were incubated in 25–50 µL of DNA extraction buffer (10 mM Tris-HCl, 2 mM EDTA, 0.2% Triton X-100, and 200 µg/mL Proteinase K in Milli-Q water) at 56 °C for 2 h to overnight. Proteinase K was inactivated at 70 °C for 10 min, and the resulting DNA solution was used as the template for PCR with KOD-FX neo (TOYOBO, KFX-201). PCR products were purified by ethanol precipitation and subjected to sequencing reactions using BigDye Terminator (Thermo, 4337456). The primers used for genotyping were as follows:

*and1*_Forward: 5′-CTCTGGCTGGTGGTACCTTG-3′

*and1*_Reverse: 5′-TTGCAGAGCTGGATCGTGTC-3′

*and1*_seq: 5′-ACAGATGCGCAGCAGCTCAG-3′

*and2*_Forward: 5′-ACAGGGCCATCTGACCAATC-3′

*and2*_Reverse: 5′-TAGCAGTCATAACCTTCCTTGG-3′

*and2*_seq: 5′-CCACAGATTGAGGACTTGGATCG-3′

PCR reaction compositions and cycling conditions followed the manufacturer’s instructions. After ethanol precipitation, the dried pellets were dissolved in Hi-Di Formamide (Thermo, 4311320), and sequencing was performed using a 3500 Genetic Analyzer (Applied Biosystems). Sequence alignment was conducted using ApE (A Plasmid Editor) with reference sequences obtained from the NCBI database and the UCSC Genome Browser.

### DAF staining and tissue fixation for zebrafish

DAF-FM DA (DAF) (Goryo Chemical, SK1004-01) staining was performed under the same conditions for all samples, regardless of subsequent experimental procedures. Living fish were incubated in 5 µM DAF solution overnight (500 µM stock in DMSO diluted 1:1000 in tank water). The following morning, fish were washed in fresh tank water for approximately 1 h. After anesthesia in 0.02% tricaine, samples (whole bodies or fin tissues) were collected, rinsed in ice-cold PBS, and fixed in 4% PFA in PBS overnight. Fixed samples were stored in PBS at 4 °C until further use. For samples not subjected to additional staining, fixed tissues were directly mounted and imaged using a confocal microscope (LSM780, Carl Zeiss) equipped with Plan Apo 10×/0.45 and C-Apochromat 40×/1.20 W objectives (Carl Zeiss). DAF fluorescence was excited and detected using a 488 nm laser line.

### DAF staining for the visualization of collagen fibers in rainbowfish fin

For whole-mount staining of rainbowfish (Melanotaenia praecox), DAF staining and fixation were performed using the same procedures as for zebrafish. For fluorescence imaging, fixed samples were mounted on glass-bottom dishes, and fin tissues were imaged using a confocal microscope (Stellaris8, Leica) equipped with 20× NA 0.8 Plan Apo objectives (Leica). Fluorescent signals from DAF-labeled collagen fibers were excited and detected using a 488 nm laser line.

### DAF staining for the visualization of collagen fibers in axolotl tissues

For whole-mount staining, living axolotl larvae at stages 47 (forelimb) and 54 (hindlimb) were incubated in 5 µM DAF solution (500 µM stock in DMSO was diluted in breeding water, 1: 1000) under dark conditions for 12 h at 20 °C. After staining, they were anesthetized with tricaine (MS-222) at an optimal concentration and fixed with 4% PFA in PBS overnight at 4°C. After the fixation, their limbs were dissected and observed with a confocal microscope (LSM780, Carl Zeiss) equipped with 20× NA 0.8 Plan Apo objective (Carl Zeiss). For the observation using a confocal microscope, the fluorescent signals of the collagen fibers stained with DAF were detected using a 488 nm laser.

### Nuclear staining of fin tissues

For Hoechst staining, fixed samples (DAF-stained at 4–5 dpf) were washed in 0.2% PBS-T at room temperature (10 min × 3) and then immersed in Hoechst 33342 solution (1:100 in 0.2% PBS-T) at 4 °C overnight. The following day, samples were rinsed with PBS and stored at 4 °C until imaging. For fluorescence imaging, samples were mounted on glass-bottom dishes, and fin tissues were imaged using a confocal microscope (LSM780, Carl Zeiss) equipped with C-Apochromat 40×/1.20 W objectives (Carl Zeiss). Hoechst fluorescence was excited and detected using a 405 nm laser line.

### Bone staining of adult zebrafish fins using Alizarin Red S

Before the experiments, the standard length (SL) of adult fish in each group was measured, and individuals with an SL of 2.1 ± 0.1 cm were selected for subsequent analyses. For Alizarin Red staining, living fish were incubated in 0.0005% Alizarin Red S solution (Sigma, A5533; diluted in tank water) overnight. The following day, fish were washed in fresh tank water for approximately 1 h. After anesthesia with 0.02% tricaine, whole fish were mounted in a glass dish, and fluorescence images of the caudal fins were acquired using a BZ-X810 microscope (KEYENCE) with an mCherry filter.

### TEM imaging for zebrafish fin tissue section

Fin tissues from larval (8dpf) and adult (with SL = 2.1 ± 0.1 cm) zebrafish were fixed overnight at 4°C in 2% glutaraldehyde (GA) and 2% paraformaldehyde (PFA) prepared in 0.1 M cacodylate buffer. Samples were post-fixed in 2% osmium tetroxide (OsO4) in 0.1 M cacodylate buffer for 2 h at 4°C, stained with 2% uranyl acetate in distilled water for 15 min at room temperature, and subsequently stained with lead solution for 3 min at room temperature. The specimens were dehydrated through a graded ethanol series (50%, 70%, 90%, and 100%), embedded in Quetol-812 epoxy resin, and polymerized at 68°C for 48 h. Ultrathin sections (70 nm) were cut using a Leica UCT ultramicrotome and examined with a JEOL JEM-1400 Plus transmission electron microscope operated at 100 kV.

### Image analysis and Statistical analysis

Imaging data were viewed and analyzed using ZEN software (version 3.5, blue edition), Imaris 10.1.1 (for cross-sectional visualization), and ImageJ (for measurements and quantification). Statistical analyses were performed using GraphPad Prism 10.

